# Temporal spiking sequences in visual cortex carry unique information about natural movies

**DOI:** 10.1101/2023.06.27.546669

**Authors:** Boris Sotomayor-Gómez, Francesco P. Battaglia, Martin Vinck

## Abstract

Information in the nervous system is encoded by the spiking patterns of large populations of neurons. The analysis of such high-dimensional data is typically restricted to simple, arbitrarily defined features like spike rates, which discards information in the temporal structure of spike trains. Here, we use a recently developed method called SpikeShip based on optimal transport theory, which captures information from all of the relative spike-timing relations among neurons. We compared spike-rate and spike-timing codes in neural ensembles from six visual areas during natural video presentations. Temporal spiking sequences conveyed substantially more information about natural movies than population spike-rate vectors, especially for larger number of neurons. As previously, shown, population rate vectors exhibited substantial drift across repetitions and between blocks. Conversely, encoding through temporal sequences was stable over time, and did not show representational drift both within and between blocks. These findings reveal a purely spike-based neural code that is based on relative spike timing relations in neural ensembles alone.

---

In recent years, the classic view on neural coding has moved from single-neuron tuning properties towards information encoding by high-dimensional populations of neurons firing patterns of spikes, which may be the basis for powerful computational mechanisms^1–8^. Yet, to make the analysis of such complex data manageable, arbitrarily defined features are typically extracted from multi-neuron time-series, potentially giving up crucial insight in favor of interpretability and simplicity of analysis. Importantly, a simple-minded and intuitive choice as firing rates, which may be combined to make up population vectors^9–15^ is likely to discard information that is used by the brain, namely the temporal structure in spike trains. Indeed, spike timing drives neural integration and synaptic plasticity^10,16–20^. Furthermore, it is unclear whether spike rates allow for stable and fast enough readout, and for sufficient coding capacity to enable cognition^17,18,21–28^. A pure spike-rate based code represents, therefore, an unwarranted simplification^12^. In fact, while it is well established that information may be extracted from spike rates after averaging, additional information can be extracted from the timing of spikes^29–36^.

Studies of temporal population coding, on the other hand, also rely on equally arbitrary, albeit intuitive, features such as the latency or phase of firing^33,37–40^. Such features often rely on a temporal reference frame only available under unnatural experimental conditions (for example the onset of a stimulus), or by assuming a regular enough neural oscillation with respect to which a “phase” may be robustly defined, an assumption often violated^41^.

As an alternative to a code based on pre-defined features, we could characterize the high-dimensional space of multi-neurons spike patterns by choosing a metric^42–45^ which attempts to capture as much as possible of the spike train’s temporal structure. Ideally, because the brain does not have access to absolute (Newtonian) time, the metric should only be based on the relative spike-timing relations among all neurons (i.e. all-to-all combinations). These relative timings are known to carry a signal that can be detected by neurons^46–48^. Here, we utilize a recently developed method called *SpikeShip*^49^ that extracts information from neural populations purely based on the relative timing of spikes alone. The method is based on solving a general optimal transport problem of determining the minimum cost of shifting ‘spike mass’ such that all relative timing relations become identical. SpikeShip designed to minimize the influence of firing rate, which allows us to study the unique contribution of temporal sequence information. Because SpikeShip is computationally efficient (it has a computational cost linear in the total number of spikes in a population), it can be readily applied to very large populations of neurons. We used SpikeShip to study the encoding of natural movies in a large population (>8000) of neurons in the visual cortex of awake mice, and contrast the information in relative timing of spikes to the firing rate. Based on the precise temporal information of spike trains, we show that multi-neuron temporal patterns convey substantially more information about natural movies than population rate vectors. Furthermore, these multi-neuron temporal patterns show high stability, i.e. no representational drift, across presentations. By contrast, firing rate codes show substantial representational drift and slowly decaying temporal correlations unrelated to stimulus content. Finally, we show that the performance advantage of temporal information increases as a function of population size and the number of active neurons.

## Results

### Comparing temporal encoding of natural movies to rate coding

SpikeShip measures the dissimilarity between spike trains of different epochs using the mathematical framework of optimal transport. It considers each spike train as a collection of “masses” (i.e. the spikes). As shown in Fig. 1A, all spikes from each active neuron, together, contribute a unit mass, which ensures the rate invariance of the method. SpikeShip solves the optimal transport problem of finding the minimum cost of shifting the (unit) mass of each neuron’s spike train in epoch *k*, such that the cross-correlations (i.e., sequential firing) between all the neurons become identical to those in another epoch *m* (See Methods). Thus, SpikeShip provides a generic dissimilarity measure of how similar all the relative spike-timing relations are between two epochs. Importantly, SpikeShip can be computed with computational cost increasingly linearly with the number of neurons (times the avg. number of spikes), and can therefore be efficiently computed for high-dimensional neural patterns. Because SpikeShip is insensitive to variation of firing rate^49^, we could determine to what extent temporal spike-timing relations carry unique information compared to firing rate population codes.

**Figure 1.**
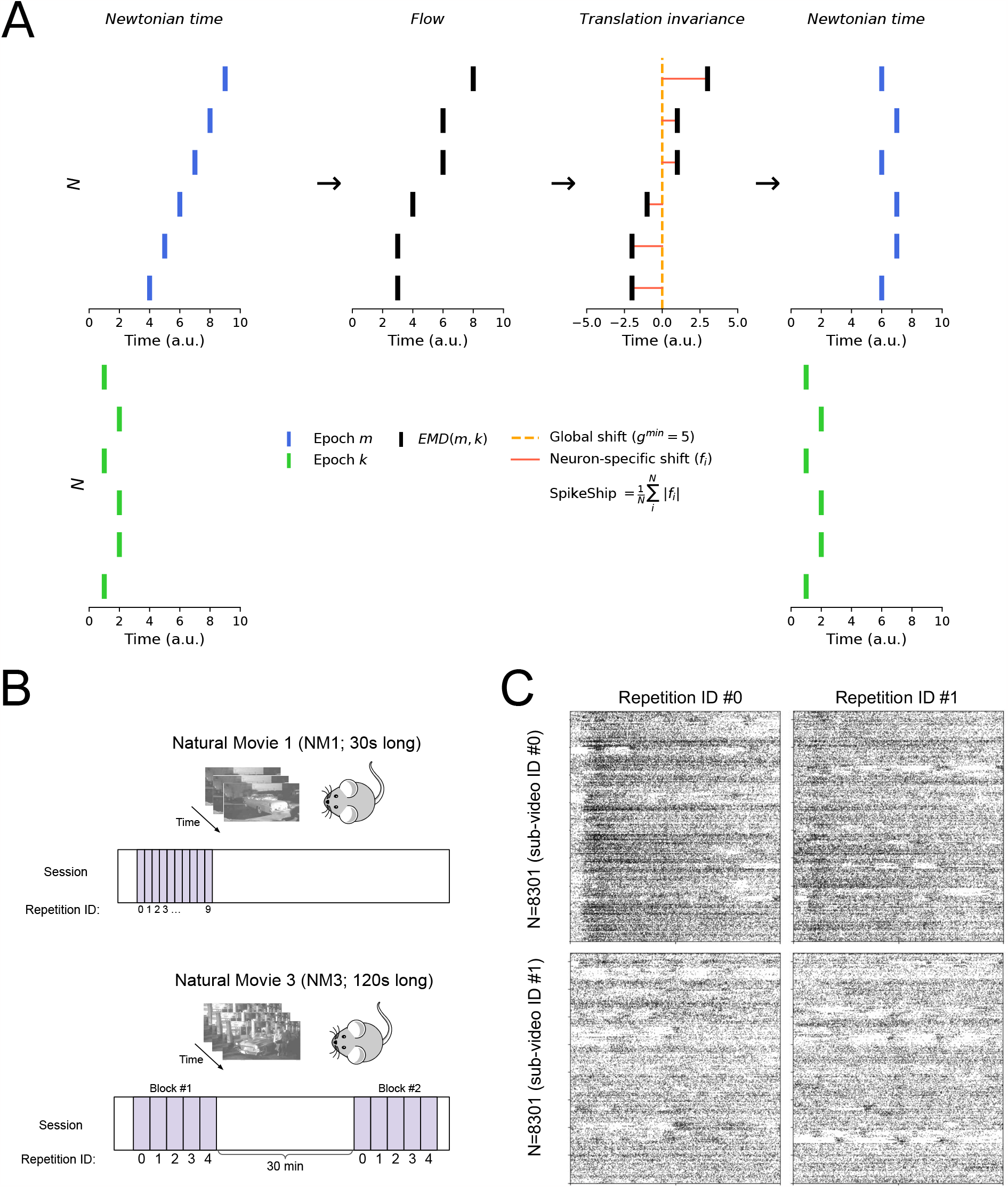
Neural population activity during natural video presentations. A) Example of SpikeShip computation of multi-neuron activity between two epochs *m* and *k*. In the first step, the difference between spike times is computed. SpikeShip correspond to the total neuro-specific flows (i.e., *f*_*i*_) relative to a global translation term (i.e., *g*^*min*^) based on minimum transport cost of spikes across neurons. SpikeShip extracts temporal information of neural sequences based on the cost to align spikes between two epochs of multi-neuronal spike trains. B) Schema of natural movies presentations from experiments from Allen Brain Institute. C) Raster plots of first (top) and second (bottom) epochs and their first (left) and second (right) repetition with *N* = 8, 301 neurons, which were pooled across 32 sessions. Each epoch/sub-video consider a window length of one second without overlap between epochs.

We analyzed Neuropixel recordings^50^ from 32 mice across six visual areas^51^ (for details, see Methods). In total, the dataset yielded population spiking patterns that consisted of *N* = 8, 301 neurons, which were pooled across multiple sessions in which the same stimuli were presented. Note that this restricts the analysis to temporal spiking patterns that are time-locked to the natural movie, however we obtained similar results when analyzing single sessions (see below). However this strategy allowed us to examine information in very large numbers of ensembles, at which the performance benefit of temporal coding was most prominent (see Fig. 1E).

For all 32 mice, we analyzed neural recordings from two natural movies: (1) *Natural Movie One* (NM1), a 30-s natural movie with 10 consecutive repetitions, and (2) *Natural Movie Three* (NM3), a 120-s natural movie presented in two blocks with 5 repetitions each (Fig. 1B). For our first set of analyses, we split the NM1 video into sub-videos of one-second length, similar to previous work^21,52^ (Fig. 1C). Thus, each epoch consisted of a dynamical stimulus of 30 frames with 10 repetitions per epoch (total *M* = 30 × 10 = 300 epochs). For each pair of epochs, we computed a dissimilarity matrix either based on SpikeShip or on firing rates (either the Euclidean distance between raw or Z-scored firing rates).

Compared to firing rates, we found that SpikeShip yielded much lower dissimilarities between repetitions of the same sub-frame as compared to pairs of different sub-frames (Fig. 2A). Correspondingly, the t-SNE embeddings of the dissimilarity matrices contained a separate cluster for each sub-video in case of SpikeShip, whereas there was substantial overlap between different sub-videos for the firing rates (Fig. 2B). Consequently, SpikeShip showed almost perfect classification of sub-videos, with much higher performance than in case of firing rates (Fig. 2C). Likewise, discriminability scores were substantially higher for SpikeShip than for firing rates (Fig. 2D). Thus, temporal spiking sequences, as quantified with SpikeShip, contained substantially more information about natural-movie content than firing rates.

**Figure 2.**
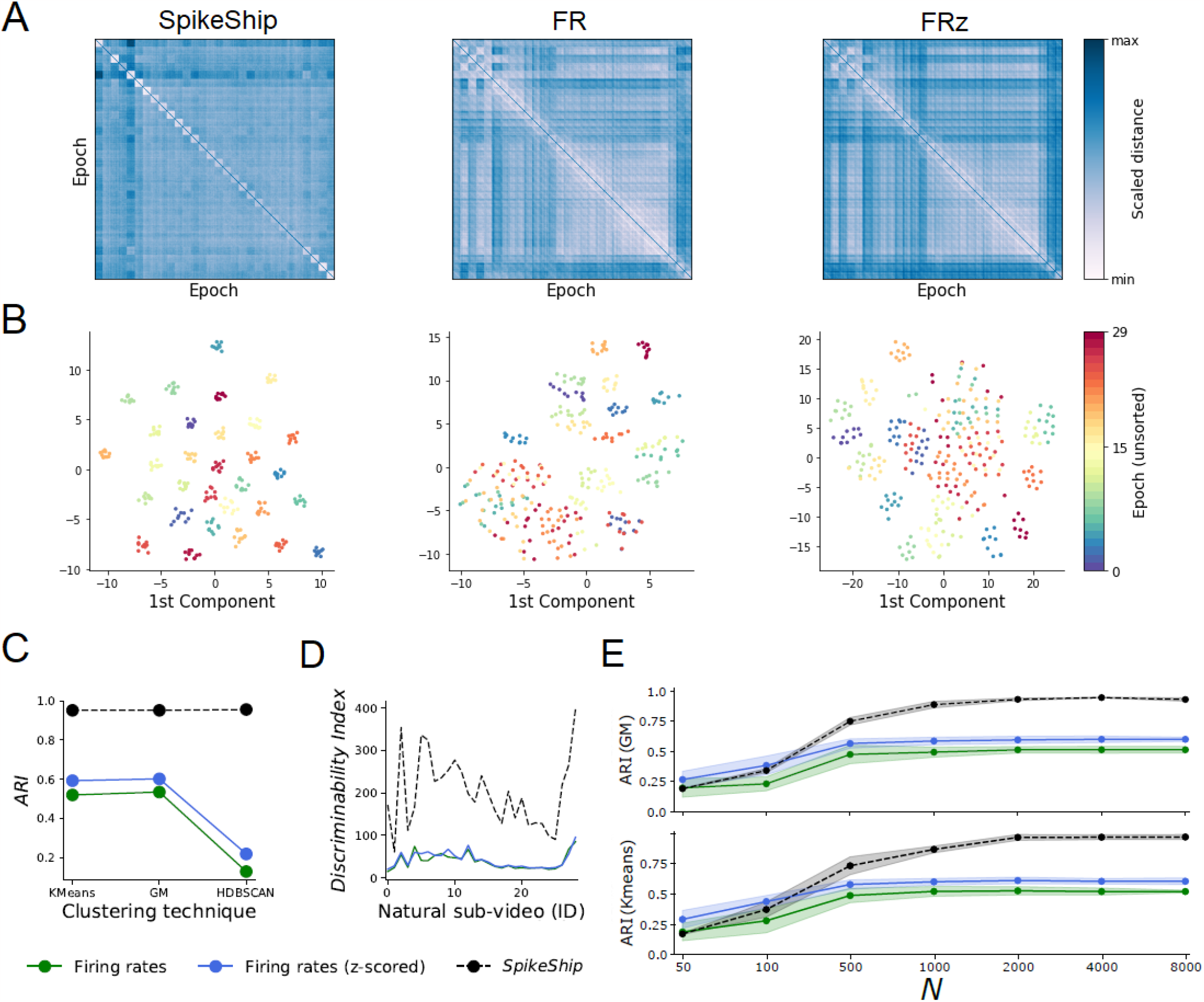
Coding of natural videos via spike count and temporal sequences. A) Dissimilarity matrices for SpikeShip, Firing rates (*FR*), and z-scored Firing rates (*FRz*). SpikeShip quantifies the similarity of epochs in terms of the relative spike-timing relationships among all neurons, based on optimal transport. Epochs are sorted by sub-video’s ID. The diagonals of dissimilarity matrices were filled with their maximum values for visualization purposes. B) 2D t-SNE embedding of pairwise distances. Colors represent one epoch (unsorted). C) Clustering performance measured via Adjusted Rand Index (ARI) using K-Means, Gaussian mixture clustering model (GM), and HDBSCAN. In the case of K-Means and GM, the number of clusters to determine in the low-dimensional embedding equals the number of distinct sub-videos (i.e., *K* = 30). D) Discriminability index for each measure across sub-videos. Discriminability indices compare the distances within clusters to those between clusters, with higher values indicating better discriminability. E) ARI Score (K-Means) by randomly subsampling *N* neurons for NM1. We performed 20 repetitions with randomly selected neurons from mice’s visual cortex. Error bars represent the standard deviation across repetitions.

We wondered whether the performance benefit of temporal sequences depended on the number of neurons that were included in the analysis. To analyze this, we took smaller subsets of neurons and repeated the clustering analysis for different subset sizes. Interestingly, the performance benefit of temporal encoding only emerged for subsets of neurons greater than about 100 neurons (Fig. 2C-E).

We also performed these analyses for single sessions (Supplementary Fig. S1A-B), which showed comparable results. SpikeShip outperformed the firing-rate code, with performance benefits emerging for population sizes of 160 neurons and beyond (Supplementary Fig. S1C). ARI scores were comparable between single sessions and pooled data across mice (Supplementary Fig. S1).

The finding that natural videos were more precisely encoded via temporal sequences than firing population vectors held true across visual areas. The best clustering performance was shown by the primary visual cortex (Supplementary Fig. S2).

To further investigate the extent to which they carry distinct information, we computed Spearman correlations between the epoch-to-epoch dissimilarity matrices of firing rates and SpikeShip (as shown in Fig. 2). These correlations can be understood as “representational similarity”. We ignored (diagonal) entries of the matrix containing dissimilarity between repetitions of the same sub-video. All the analyses were performed using Mantel-test^53^ to compute correlations between two dissimilarity matrices. SpikeShip and firing rates a moderate (Spearman) correlation around 0.26 (*p* ≤ 0.001), demonstrating largely distinct information in spike counts and temporal sequences.

The latter result held true across different visual areas (Supplementary Fig. S3), with Spearman correlations between 0.02 and 0.29. We also performed representational similarity analyses by comparing the dissimilarity matrices between different visual areas. Interestingly, we found that representational similarity between areas was stronger for firing rates than for SpikeShip (Supplementary Fig. S4). In other words, different visual areas provided more independent information through temporal sequences than through the population rate vectors.

### Correlations in neural representations on short time scales

In Fig. 2, we had computed the dissimilarity matrix between all pairs of epochs. We wondered how neural representations varied during the progression of the movie, e.g. whether epochs that were closer in time within the movie have similar representations. Indeed, the t-SNE representations for firing rates suggest that epochs that are close in time lie closer to each in the low-dimensional embedding space (Fig. 3A). By contrast, a similar effect was not observed for SpikeShip, with an apparent random ordering of clusters w.r.t. time.

**Figure 3.**
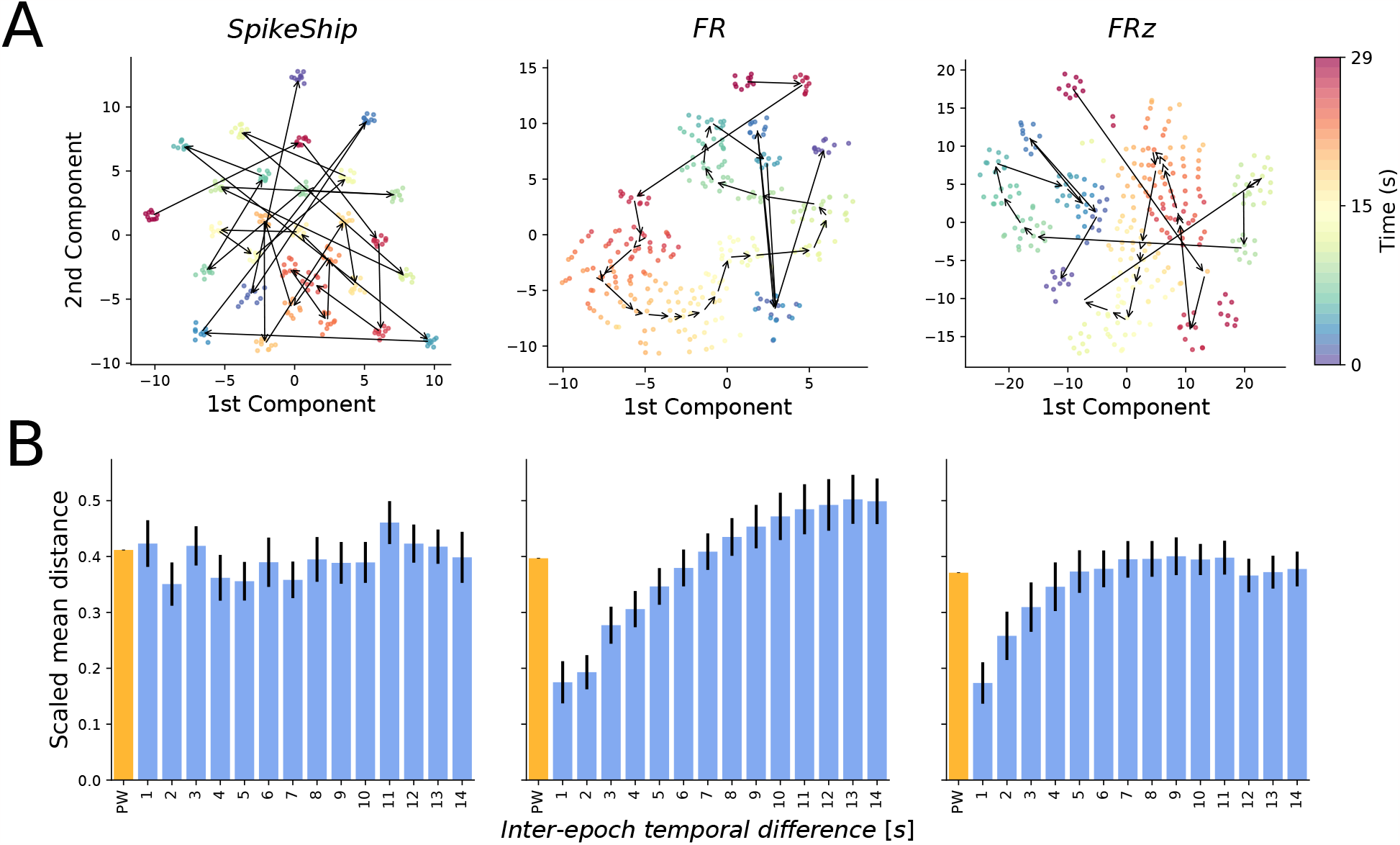
Rate-based neural activity reflect ordering in natural movies. A) Relation of neural activity to elapsed time in natural movie. Color represents the time of epoch presentation (i.e., second of the movie). Arrows represent the direction of the trajectory between consecutive sub-videos (centroids). Population firing rates form a continuous trajectory, whereas temporal sequence representations are discontinuous. B) Mean scaled (Euclidean) distances across full pairwise distances between epochs (PW) and consecutive epochs. Numbers indicate the time (in seconds) between two consecutive epochs. The black line represents the standard deviation divided by the square root of the number of samples (i.e., 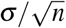).

To quantify this, we computed the Euclidean distances between consecutive epochs and compared these with distances to other epochs (to compare the different measures, these distances were computed in t-SNE space). For firing rates, the distance between two consecutive epochs is about 3-fold lower than the distances between all epochs (i.e. pairwise comparison) (Fig. 3B). This resulted in correlated firing rate patterns up to about 7 movie epochs (Fig. 3B). By contrast, for SpikeShip we did not find a difference between consecutive epoch pairs and random epoch pairs.

Additionally, we examined the autocorrelation of mean firing rate vectors across sub-videos of NM1. We observed a high correlation between consecutive sub-videos (Pearson product-moment correlation index, Fig. S5). The correlation value decay up to about 10 movie epochs. Together these findings suggests that firing rate representations tend to be similar for nearby epochs, indicating correlations on the time scale of seconds. This causes a smearing that reduces the information content about the movie. By contrast, temporal patterns show uncorrelated representations across movie epochs, resulting in highly precise encoding.

### Representational drift of firing rate and temporal code

Next, we investigated to what extent the coding of natural movies via spike rates or temporal sequences was stable within a session. Several studies have reported that neural representations of stimuli, tasks or contexts can change across time within a session^21,22,54^.

To examine this, we computed the correlation of neural activity across different repetitions of the same sub-video. The low-dimensional embedding colored by the repetition order shows ample variability across repetitions for *FR* and *FRz*, and little variability for SpikeShip (Fig. 4A). In particular, firing rates map out a trajectory in the low-dimensional embedding space during each repetition, and this trajectory gradually drifts away across repetitions. To further quantify this, we computed for each repetition a vector of Eucldean distances across the sub-videos, and then correlated these vectors between repetitions. This yields a matrix of correlations ordered by repetition number (Fig. 4B). The population vector of firing rates decorrelated gradually across time, as shown in Fig. 4C. Yet, for SpikeShip, the Pearson correlation stayed close to 1 for all time-delays. Similar findings were made by analyzing the distance between dissimilarity matrices (Fig. S6).

**Figure 4.**
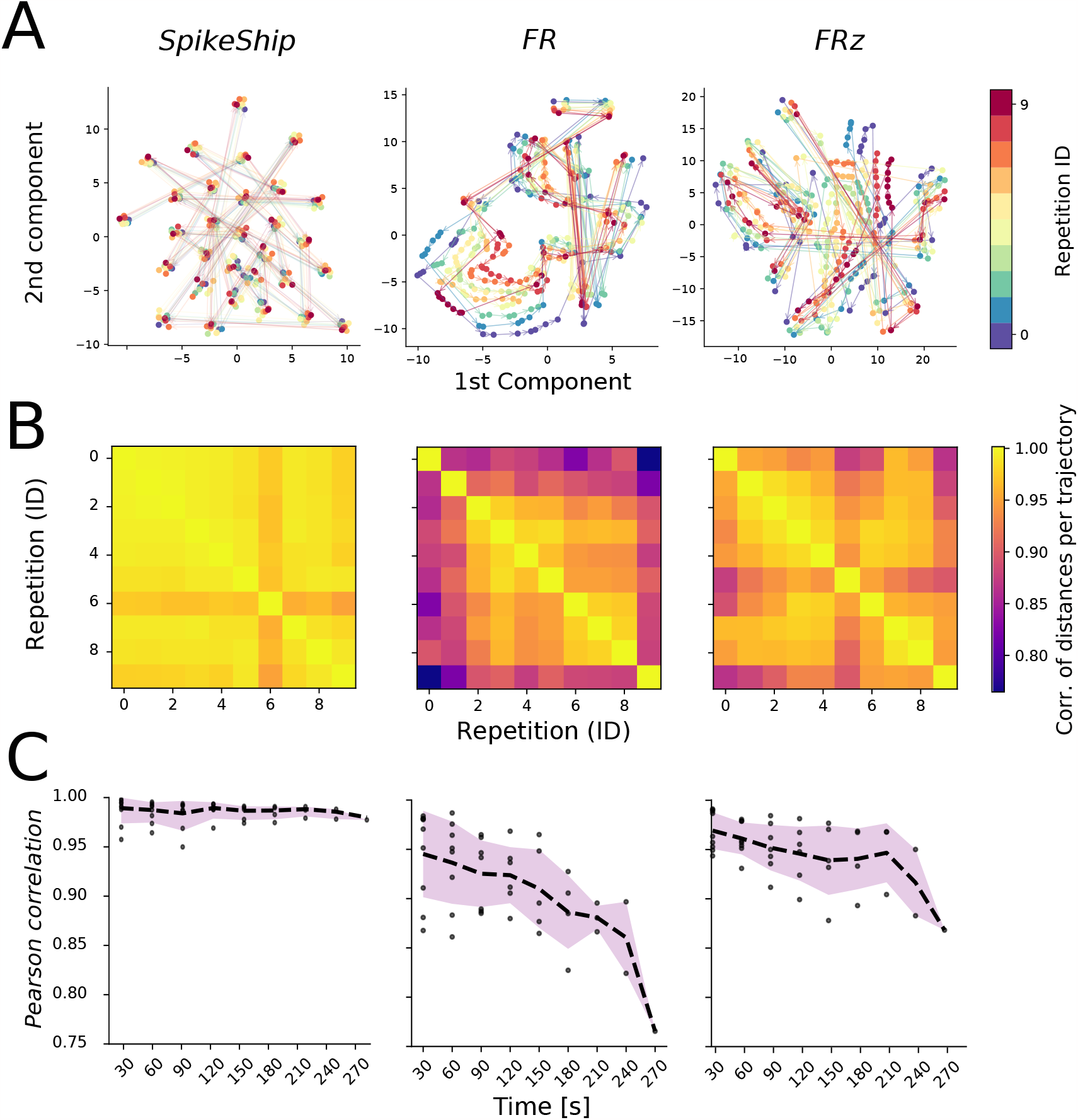
Changes in neural representations across repetitions. A) 2D t-SNE embedding of pairwise distances. Color represents the repetition ID of each Natural Movie presentation. Arrows represent the direction of the trajectory between consecutive sub-videos. Each trajectory corresponds to one repetition. B) For each repetition, the Euclidean distance was computed between sub-videos. Pearson correlations were then computed between these vectors of Euclidean distances. C) Pearson correlation as a function of time.

Together, these analyses indicate that the temporal structure in the spike trains (quantified with SpikeShip) uniquely and reliably encodes the different dynamic stimuli, with essentially no drift in the population code. The firing rate on the other hand showed systematic drift within a session.

To generalize these results, we analyzed another dataset (NM3) in which longer movies were presented (5 repetitions), repeated in two separate blocks with 30 minutes between blocks. Similar to the analyses presented above, we found a tight clustering of the different sub-videos for SpikeShip (Fig. 5A), while different subvideos were not well separated for firing rates (Fig. 5A). To quantify drift, we again computed Pearson correlations between the vectors of Euclidean distances. Firing rate representations showed drift both within the blocks (across repetitions) and between blocks (Fig. 5D). SpikeShip, however, showed Pearson correlations close to 1 indicating stable encoding of natural movies based on temporal sequences (Fig. 5D).

**Figure 5.**
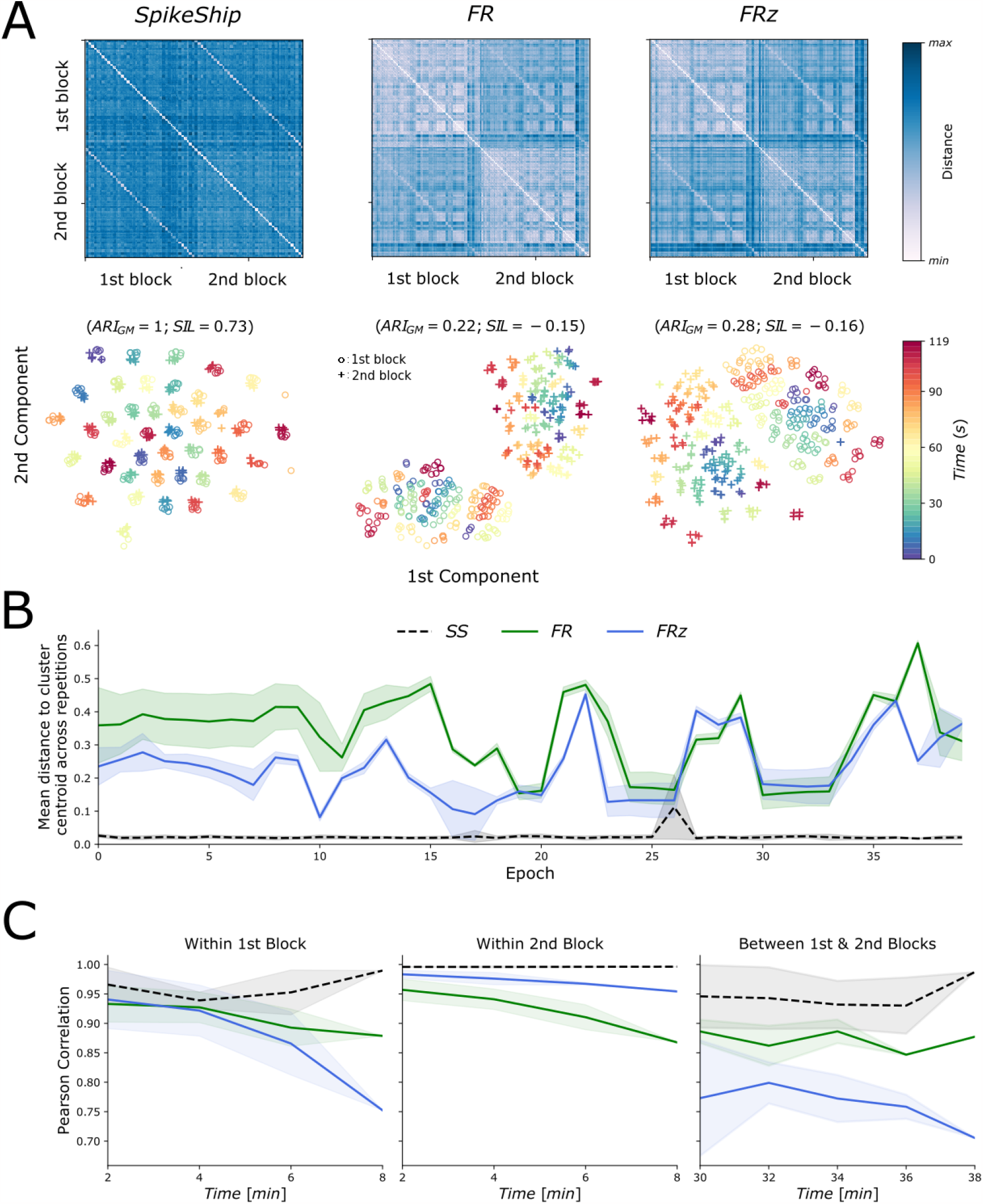
Rate-coding vs temporal coding for long natural movie presentations (NM3). A) Dissimilarity matrix for Firing rates (*FR*), z-scored Firing rates (*FRz*), and SpikeShip. Epochs are sorted by sub-video’s ID and block. B) 2D t-SNE embedding of pairwise distances. Colors represent sub-video’s start time. C) Mean distance to cluster centroids across repetitions. D) Correlation analysis within and between blocks of NM3 presentations, similar to Fig. 4C.

### Dependence of encoding on number of active neurons

Finally, we investigated to what extent the performance of SpikeShip depended on the number of active neurons. For example, it is possible that the advantage of SpikeShip vs. firing rates is particularly prominent during epochs in which there are few active neurons, or alternatively during epochs in which there are many active neurons. To this end, we quantified for each sub-video of 3 frames the number of active neurons. The number of active neurons was related to stimulus content. Specfically, there was a strong correlation between the number of active neurons and the curvature across frames^55,56^ which measures the variability of movie’s pixel across frames (*Global Curvature* (GC)) (Fig. 6A-B). We wondered how the clustering performance of firing rates and SpikeShip depended on the number of active neurons for these shorter sub-videos. We selected epochs that contained either the highest or the lowest number of active neurons for a given pair of epochs. When the number of active neurons was high, SpikeShip yielded strong clustering even for short sub-videos (Fig. 6C). By contrast, when the number of active neurons was low, SpikeShip did not yield clear clustering, whereas firing rates yielded high ARI scores (Fig. 6D). Hence, temporal coding, as quantified with SpikeShip, is the most reliable coding scheme when the number of active neurons is high, whereas firing rates are more reliable when the number of active neurons is low.

**Figure 6.**
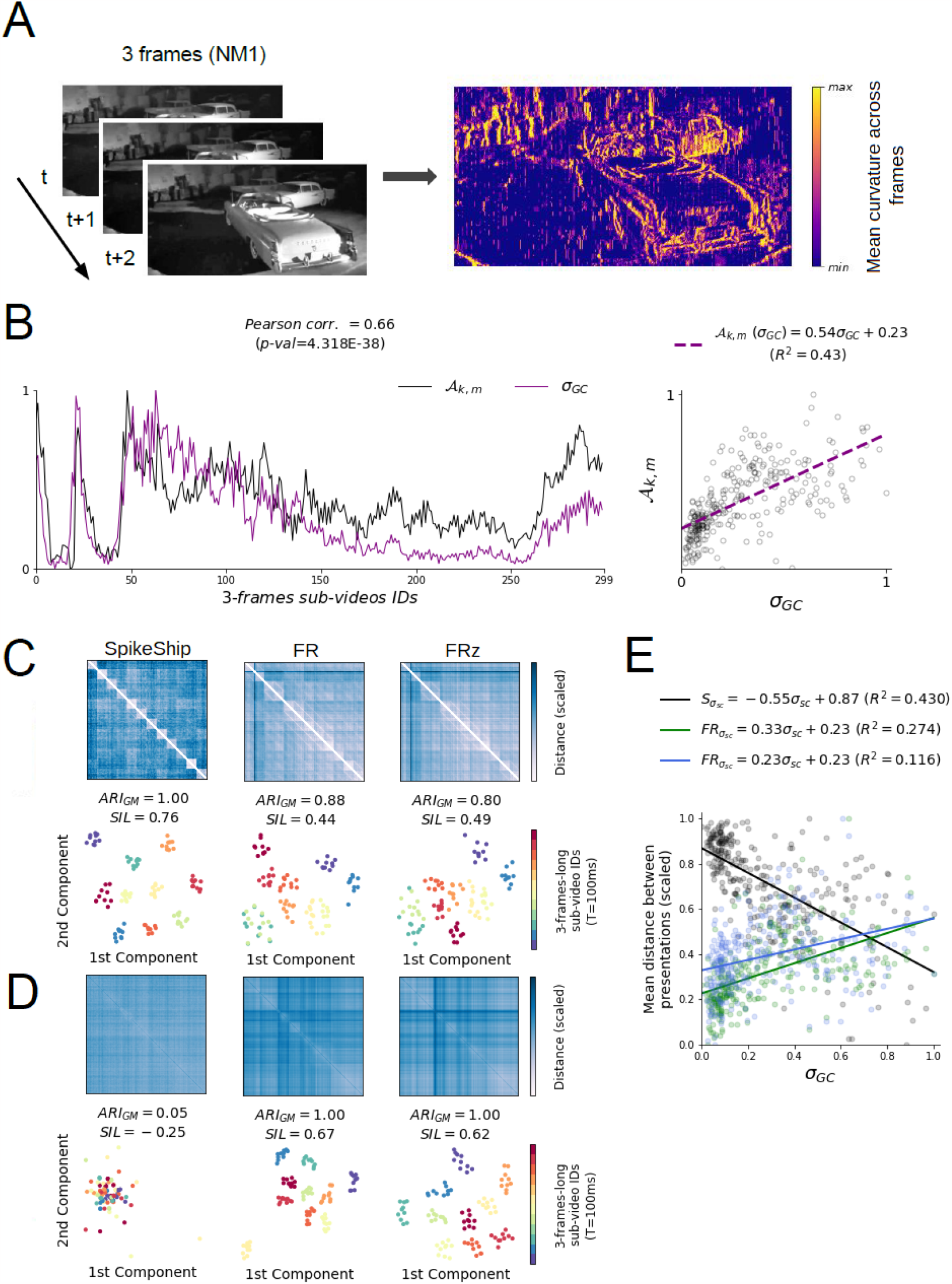
Relationship of population coding to number of active neurons. A) Summary of computation of mean curvature across sub-videos of 3-frames (*≈* 100ms). Pixel distance of natural movies frame is computed as in^55,56^, mentioned as Global Curvature (GC). B) Raster plot of the proportion of active neurons between all the pairs of epochs *k* and *m* (*𝒜*_*k,m*_) and the standard deviation of the global curvature across frames (*σ*_*GC*_). Pixel variability *σ*_*GC*_ across sub-videos is correlated with the rate of active neurons *𝒜*_*k,m*_ (i.e., Pearson correlation = 0.66). C) 10 epochs with the highest variability of the global curvature (*σ*_*GC*_). Top: Dissimilarity matrices for firing rates (FR), z-scored firing rates (FRz, normalization across epochs), and SpikeShip (S). Bottom: 2D t-SNE embeddings with precomputed distance matrix. To measure clustering performance we used *ARI*_*GM*_ and *SIL* which represent Adjusted Rand Index using Gaussian Mixtures and Silhouette, respectively. D) 10 epochs with the lowest variability of the global curvature (*σ*_*GC*_). Top: Dissimilarity matrices for firing rates, z-scored firing rates (normalization across epochs), and SpikeShip. Bottom: 2D t-SNE embeddings with precomputed distance matrix. E) Multi-neuron distance depends on *σ*_*GC*_. Pearson correlation coefficients of 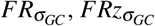, and 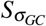 are 0.52 (p-val *<* 10^*−*220^), 0.34 (p-val *<* 10^*−*90^), and *−*0.66 (p-val *<* 10^*−*381^), respectively.

To further quantify this, we correlated the mean dissimilarity between different presentations of the same sub-video as a function of the global curvature, which is a proxy of the number of active neurons. We found a negative correlation for SpikeShip, i.e. representations tended to be more similar (i.e. less precise coding) when then global curvature was high. By contrast, for firing rates we found a positive correlation, i.e. representations tended to be more dissimilar (i.e. more precise coding) when the global curvature was high.

## Discussion

We investigated how large populations of neurons in visual cortex encode natural videos. Instead of extracting features from the neurons’ spike trains like firing rate or latency, we took an approach to neural coding based on optimal transport theory without pre-defining features. Specifically, we used SpikeShip, a recently developed method that allows for unsupervised detection of multi-neuron patterns exclusively based on the relative spike-timing relations among neurons. Due to its linear computational cost, we could compute SpikeShip up to several thousands of neurons. Strikingly, we found that natural videos were encoded much more precisely through the relative spike-timing relations as compared to the firing rate. Furthermore, the encoding of natural videos through spike timing was highly stable across time, whereas firing rate population vectors showed substantial representational drift within sessions. Temporal coding had strong performance advantages for large neural populations, and during epochs in which many neurons were active, while rate codes showed better performance during epochs of sparse firing, indicating that both codes may provide synergistic and complementary information.

The present study suggests that inter-neuronal spike-timing relations encode natural-video content with high precision. Importantly, we demonstrated this result with SpikeShip, which is a measure in which the energy of each spike train is normalized. That is, SpikeShip is invariant to the total spike count in a temporal window. We found that population firing rate vectors were less reliable than SpikeShip in encoding information. These results raise the question of how downstream post-synaptic neurons can read out the information encoded by the pattern of relative spike-timing relations. This would require that single neurons discriminate different patterns of synaptic inputs based on their relative arrival time. Experimental evidence shows that the impact of a sequence of synaptic inputs does indeed depend on the relative timing between the synaptic inputs^48^. Such sensitivity to specific sequences likely depends on dendritic non-linearities. The recurrent interactions among neurons in a post-synaptic population may further facilitate the read-out of specific sequences: Recent work suggests that the read-out of specific temporal sequences is enabled by a combination of spike-time-dependent-plasticity and inhibitory feedback^57,58^. Importantly, our theoretical work suggests that the computational load of reading out temporal sequences is in theory extremely low, and equal to the spike count^49^. This would suggest that relatively efficient read-out mechanisms could exist that are implemented via recurrent neural networks.

By disentangling relative spike timings and firing rates with SpikeShip we could examine the drift in firing rates and relative timing relations between neurons, quantified with rate-invariant measure. Firing rate drift does align with previous research on drift in representations^21–23,28,59–63^. By contrast, we found that there was essentially no drift in the relative spike-timing relationships among neurons, i.e. in the sequential firing of neurons. This finding may shed light on the function and mechanisms of representational drift. There are opposing mechanistic accounts of representational drift. Some studies suggest that changes in behavioral state contribute to drift^52,64^, while others propose changes in the synaptic weight matrix^65,66^. It remains to be investigated why these factors would impact population rate vectors, but not the encoding of natural videos via the temporal structure of spike trains.

The present study matches will with several previous studies comparing the encoding of natural videos between spike count and temporal coding.^67^’s study in anaesthetized monkey V1 agrees with ours, by demonstrating additional information through relative timing compared to firing rates. However, the present study addressed this by analyzing relative spike-timing in high-dimensional neural ensembles, whereas^67^ focused on individual neurons and the encoding of information through the spike-LFP phase. It remains to be investigated to what extent fluctuations in spike-LFP phase can be translated into the relative spike-timing relations among large numbers of neurons. Importantly, the present study also suggests that the power of temporal coding becomes apparent only for large ensembles of neurons, in which a sufficient number of cells fires a spike.

Our finding that the relative spike-timing relations encoded information with similar precision when pooling neurons indicates that these spike-timing relations resulted from the temporal structure of the video itself. In this sense, the sequential firing of neurons differs from e.g. intrinsic sequences encoding information via theta sequences in the rodent hippocampus^68–72^, or via gamma phase shifting in visual cortex^73^. It remains unknown to what extent the temporal encoding of natural video results from feedforward activation, or in addition from recurrent neural interactions. However, it is likely that the temporal relations result from a combination of both, i.e. feedforward drive resulting in specific temporal patterns via recurrent interactions among neurons^33^.

Together, our findings offer a new view on neural coding that is based on the temporal structure of spike trains and which can be captured by a method based on optimal transport theory that has the same computational complexity as the spike count. The ensuing coding strategy is purely spike-based and does not presuppose a notion of firing rate. Together, our findings suggest that information about natural visual input is encoded robustly and stably by high-dimensional temporal spiking sequences.

## Methods

### Natural movie’s processing from Allen Brain Institute datasets

We used the public available datasets of Allen Brain Institute through AllenSDK (For more details, see http://help.brain-map.org/display/observatory/Documentation). Neuropixels silicon probes^50^ were used to record neurons with precise spatial and temporal resolution^51^. We selected the cells of 32 mice during natural scenes presentations. The cells were selected considering a signal-noise ratio (*SNR*) such that *SNR* > 0. The neural activity from a total of *N* = 8, 301 cells was selected from the Primary visual area (VISp), Lateral visual area (VISl), Anterolateral visual area (VISal), Posteromedial visual area (VISpm), Rostrolateral visual area (VISrl), and Anteromedial visual area (VISam).

Finally, for the analyses per brain area we present in Fig. S2, we down-sampled the set of neurons randomly in order to compare the performance of both decoding schemes. Particularly, we used *N*_*ds*_ = 879 for every brain area.

### Computation of dissimilarity matrices of firing rate vectors

For a population of *N* neurons, we computed the firing rate vectors 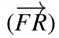 for each epoch of our analyses as the count of spikes per neuron divided by a window length *T*. We denote *FRz* the normalized firing rate vectors across epochs (z-score). Finally, we computed the Euclidean distance between both normalized vectors for two epochs *m* and *k* as 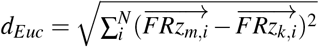.

### Computation of dissimilarity matrices via Spike-Ship

SpikeShip is a dissimilarity measure based on optimal transport theory to extract temporal multi-neuron spike-train patterns. SpikeShip solves the following transport problem:

Suppose a population of neurons in two epochs *k* and *m*. For each neuron *j* in epoch *k* for which the number of spikes *nk, j* > 0, we define the point process with unit energy

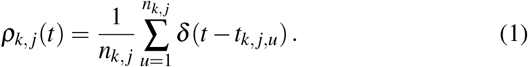

r
This defines for each pair of neurons (*i, j*) in epoch *k* the cross-correlation function

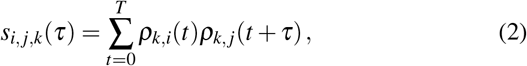

Consider two epochs (*k, m*). We wish to find for each neuron (in epoch *k*) a transport of mass from *t* to *t*′, [**M**] _*j,t,t*′_, such that *s*_*i, j,k*_(*τ*) = *s*_*i, j,m*_(*τ*) for all (*i, j, τ*). The mass here consists of the spikes, which have a sum of 1. The objective is then to find a matrix of flows **M** that minimizes the total mover cost, i.e.

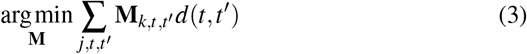

where *d*(*t, t*^*′*^) = |_*t*_ *−*_*t*_^*′*^| (For more details, see^49^).

### Discriminability index

We quantified these differences within and between natural images for each stimulus *id* in a matrix labelled as *diss*.

To this end, the calculated a “Discriminability index” (*d*) defined as the disparity between the average distances within and between stimulus *id*, divided by the square of the sum of their variances:

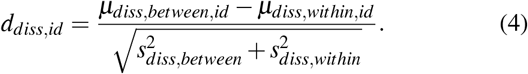

This index indicates how many standard deviations are two sets of distances away from each other (For details, see our previous work^49^).

### Measure of representational drift

#### From 2D t-SNE embeddings

For each dissimilarity matrix of NM1 and NM3 presentations, we computed the Euclidean distances across consecutive epochs (i.e., trajectory). Then, we measure the representational drift as the Pearson rank correlation between pairs of trajectories, as done in^21^. Such correlations are visualized across the time difference between repetitions of the same movie.

#### From dissimilarity matrices

We extracted the upper triangle based on the pairwise distances of each scaled dissimilarity matrix (Fig. S6A). For a dissimilarity matrix *raw*_*diss*_*m*_ based on metric *m* with *m ∈ {FR, FRz, S}*, we compute:

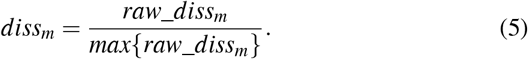

Finally, we computed the Euclidean distance between dissimilarity matrices from each pair of repetitions, as shown in (Fig. S6B).

### Global curvature definition

In order to compute the changes in the pixel domain of the movie, we based on previous studies about curvature computation of pixels in movies^55,56^. We denoted *x*_*t*_ to one pixel of the movie from the frame *t*. First, we computed the difference of such pixel at time *t* with the previous one (*t −* 1) as

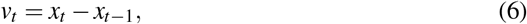

as shown in Fig. 6A).

We normalized such vector based on its norm as:

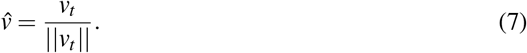

Then, the local curvature is defined as

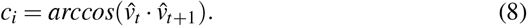

This measure is always a positive number. It reaches its lowest value of zero only when the frames are in a straight line without any bending in the high-dimensional space. Thus, the global curvature correspond to the average of all local curvatures over time, computed as

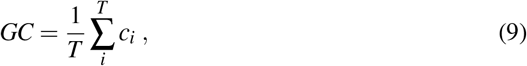

in degrees. See Henaff et al. studies^55,56^ for further details.

## Acknowledgements

This project was financed by the BMF (Bundesministerium fuer Bildung und Forschung) to MV, Computational Life Sciences, project BINDA (031L0167) to MV, an ERC starting grant (850861) to MV, DFG VI Grants (908/5-1 and 908/7-1) to MV, and the NWO VIDI Grant to MV.

## Author contributions statement

Conception and problem statement: BSG, MV. Data analysis and simulations: BSG. Mathematical analysis: BSG and MV. Supervision: MV, FB. Writing of main draft: BSG, FB and MV.

## Supplementary information

**Figure S1.**
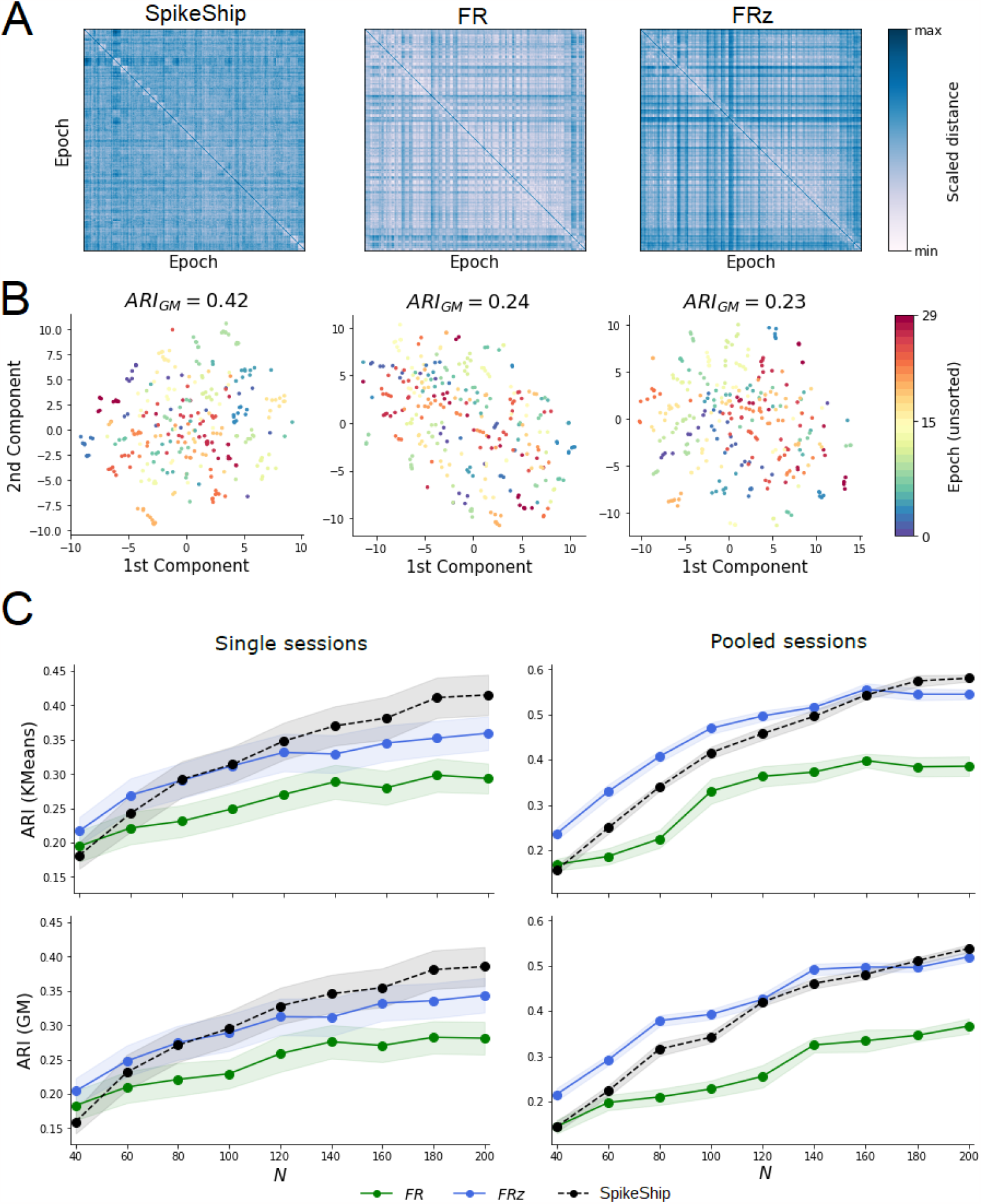
Population coding of natural videos for single sessions. A) Dissimilarity matrices sorted by sub-video presentation time. *M* = 300 epochs (30 sub-videos with 10 repetitions each) and *N* = 365 neurons. The diagonals of each dissimilarity matrix contain the maximum values for visualization purposes. B) 2D t-SNE embeddings from pre-computed values shown in A). Color represent the sub-videos (unsorted). C) Clustering performance for an increasing number of neurons across single sessions (Left) and pooled sessions (Right). We selected sessions with minimally 200 neurons. *ARI*(*KMeans*) and *ARI*(*GM*) correspond to the Adjusted Rand Index using KMeans and Gaussian Mixtures (GM) as clustering techniques, respectively. Error bars correspond to the standard deviation across sessions divided by the square root of the number of sessions (i.e. 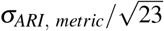).

**Figure S2.**
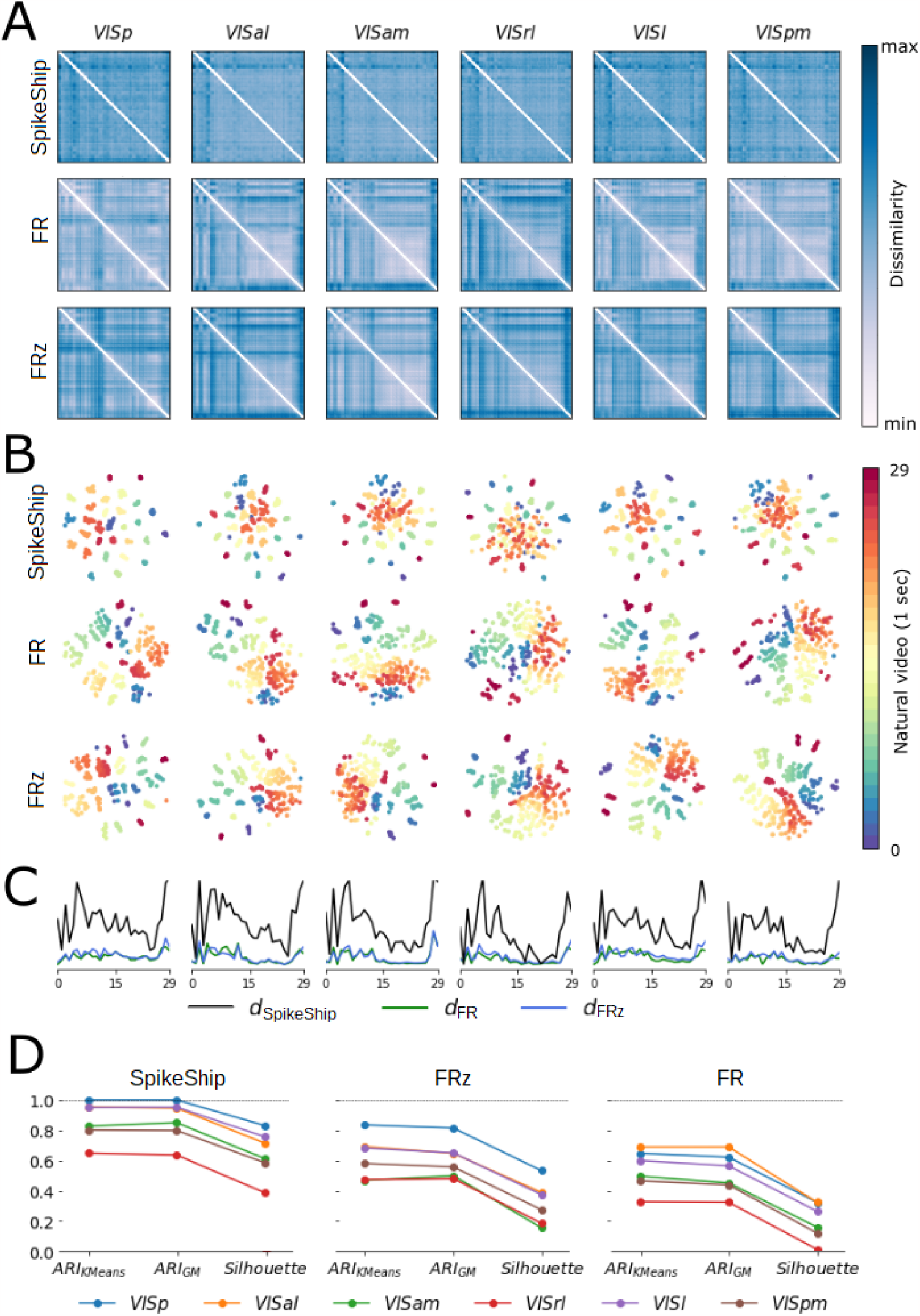
Reliability of time-coding scheme is preserved across brains areas. A) Dissimilarity matrices per brain area (columns) and measures (rows). B) 2D t-SNE embeddings per brain area (columns) and measures (rows). C) Discriminability index for each measure across sub-videos and brain areas. D) ARI and Silhouette score for each measure across sub-videos and brain areas.

**Figure S3.**
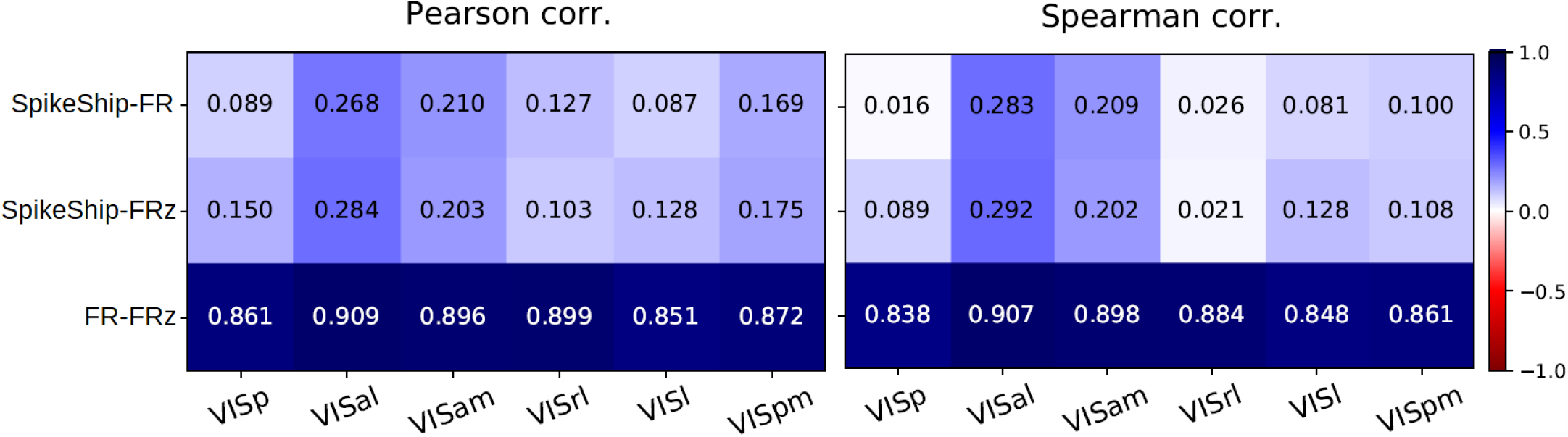
Representational similarity analysis between firing rate and SpikeShip, similar to Fig. 3, but now for separate areas. Left: Pearson correlation coefficient, Right: Spearman correlation coefficient. Both coefficients show that the relation (SpikeShip, *FR*) and (SpikeShip, *FRz*) are close to zero across visual brain areas.

**Figure S4.**
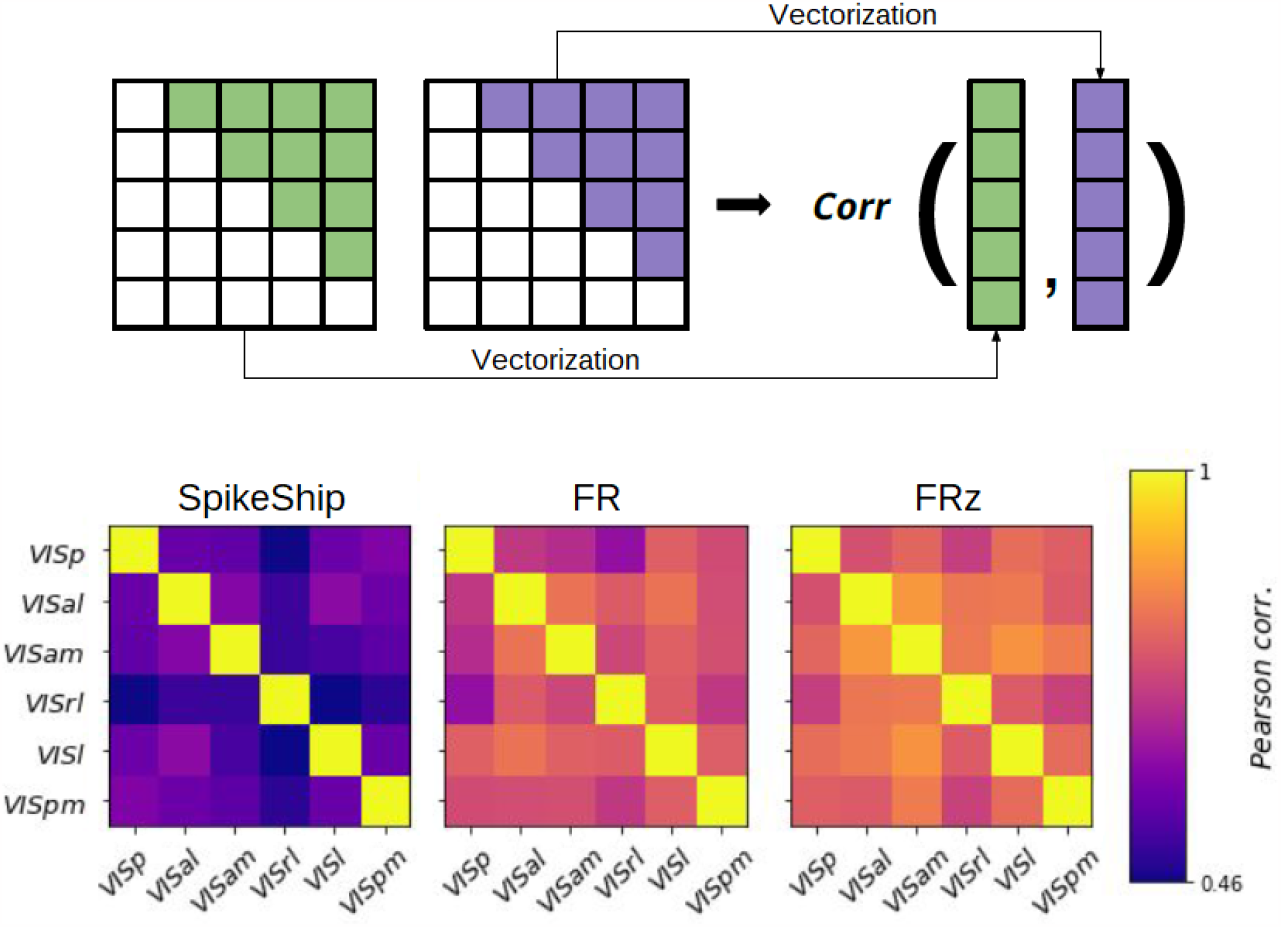
Representational dissimilarity analysis. Top: Summary of vectorization (representational vectors) process for dissimilarity matrices and computation of pairwise correlations. Bottom: Pairwise correlation of representational vectors using Pearson correlation coefficient per visual area across 32 mice (*N* = 8301).

**Figure S5.**
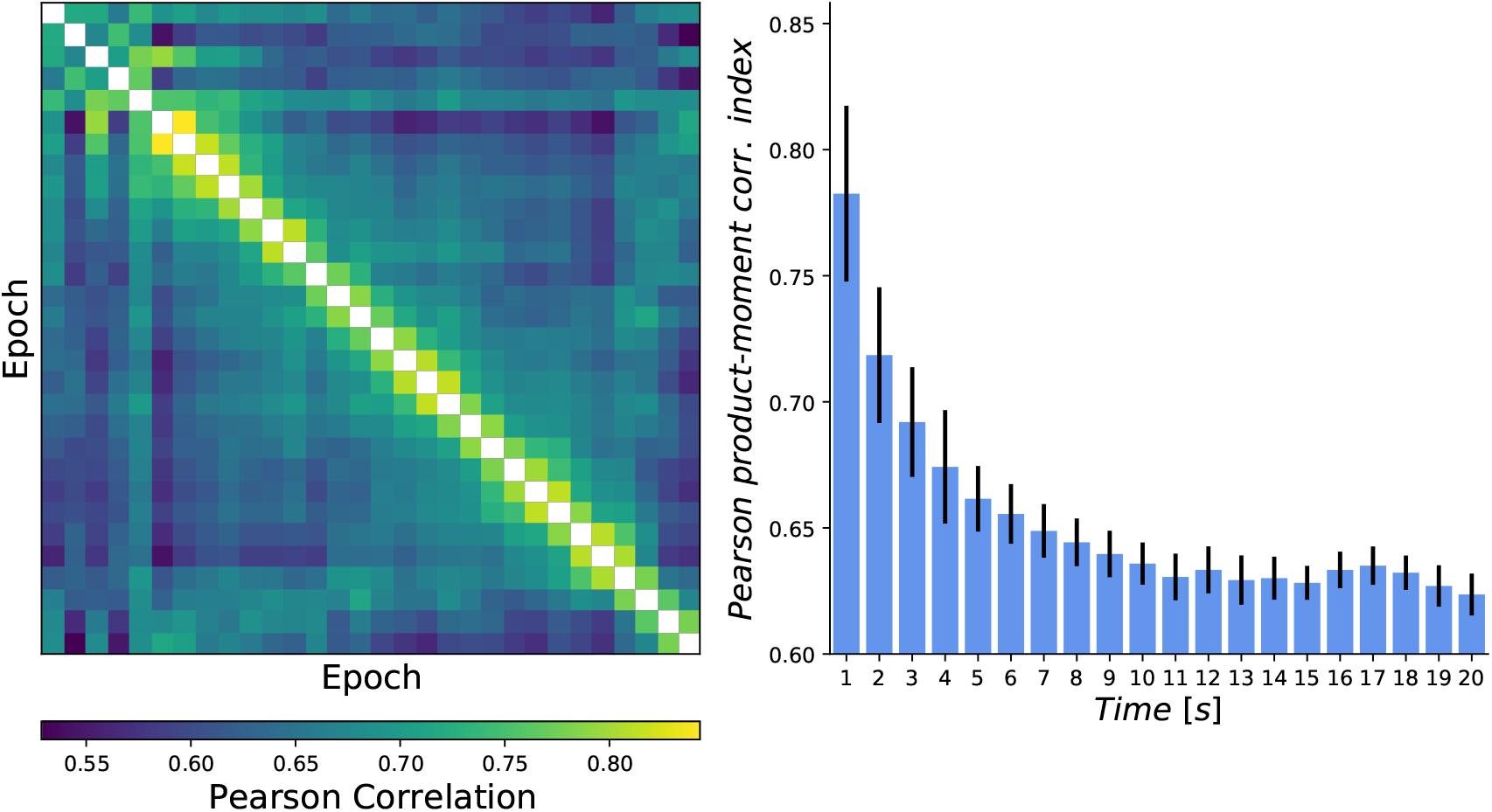
Correlation of mean spike counts vectors across sub-videos of NM1. Left: Mean Pearson product-moment correlation index between pairs of epochs. Right: Mean correlation between epochs with different separations in time. Error bars correspond to the standard deviation across sessions divided by the square root of the number of epochs.

**Figure S6.**
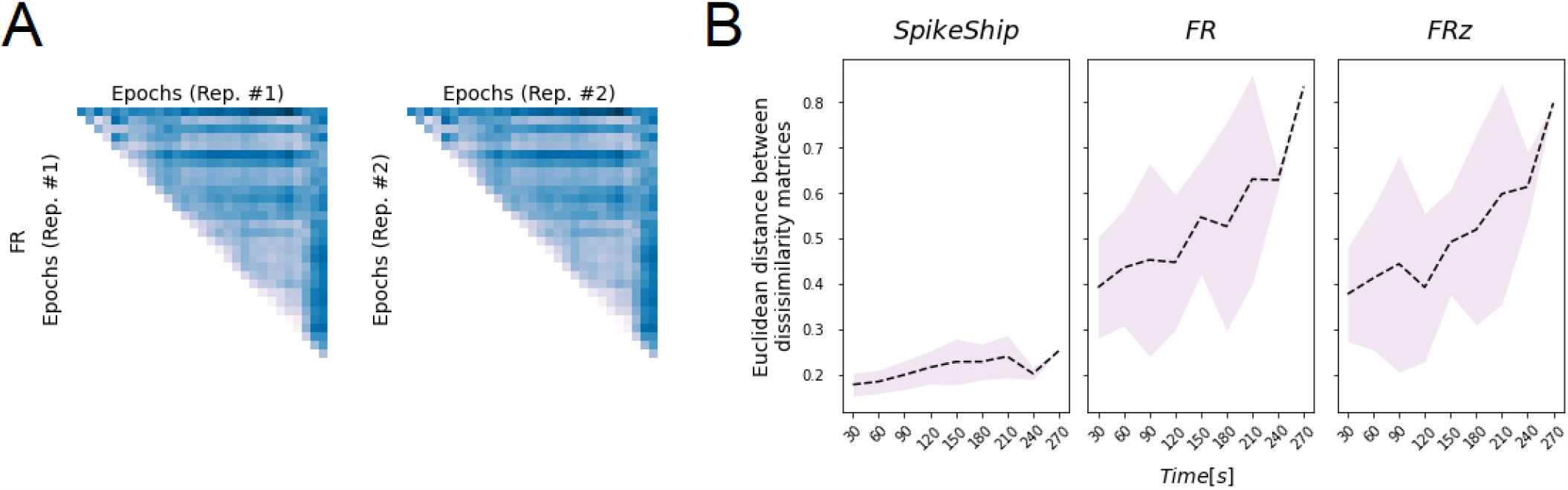
Representational drift as the distance between dissimilarity matrices. A) Example of dissimilarity matrices for two repetitions. Each matrix contains the dissimilarity of multi-neuron spike sequences across the entire movie (M=30 epochs, with sub-videos of 1 second). Epochs are sorted by time, as in^21^. B) Euclidean distance between dissimilarity matrices in function of the time difference between two repetitions. Lines and filled regions represent the mean and standard deviation of the correlation for each metric.

**Figure S7.**
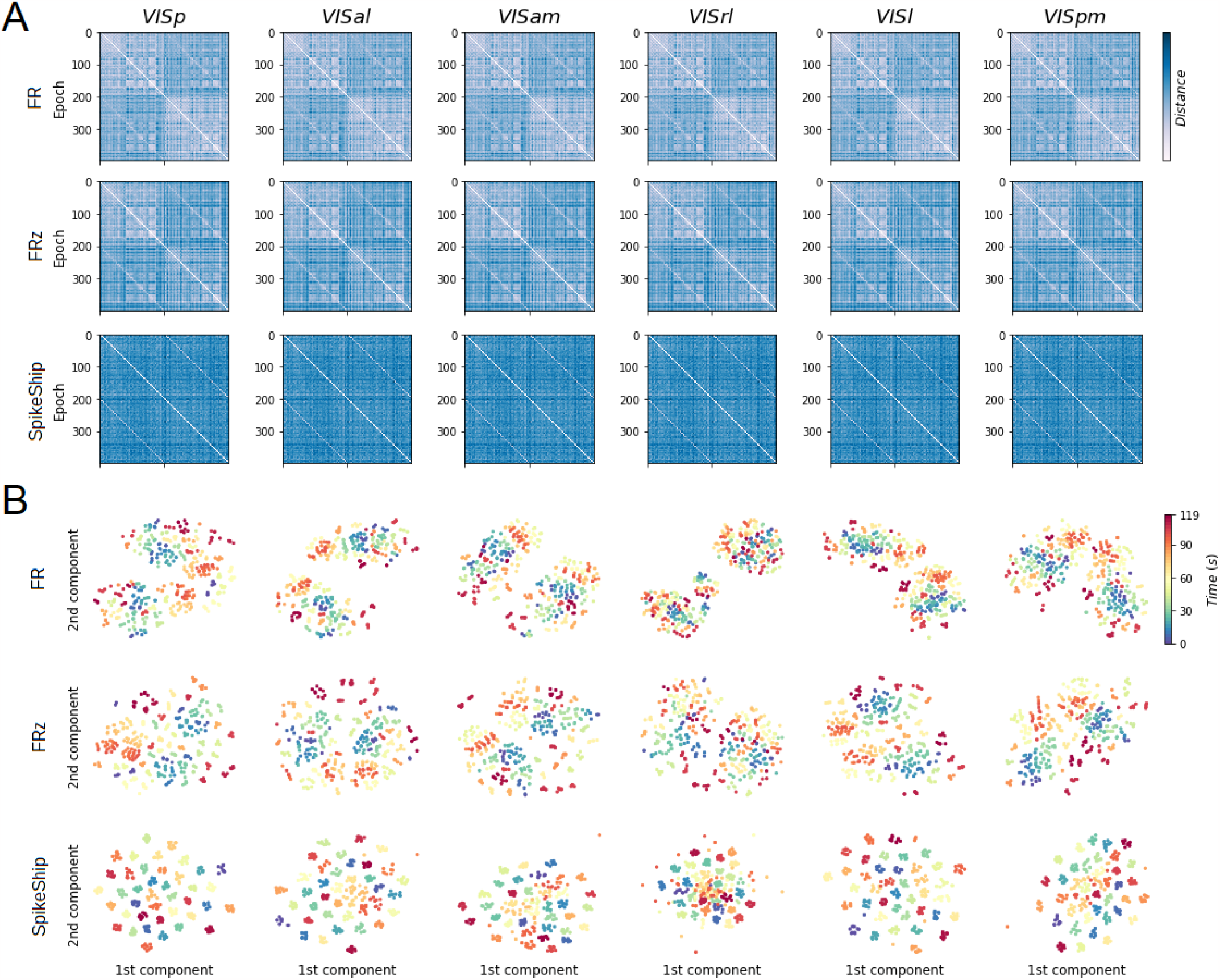
Reliability of time-coding scheme is preserved across brains areas and blocks of natural movie presentations. A) Dissimilarity matrices per brain area (columns) and measures (rows). B) 2D t-SNE embeddings per brain area (columns) and measures (rows).

